# Genetic control of *Campylobacter* colonisation in broiler chickens: genomic and transcriptomic characterisation

**DOI:** 10.1101/2020.04.14.040832

**Authors:** Androniki Psifidi, Andreas Kranis, Lisa M. Rothwell, Abi Bremner, Kay Russell, Diego Robledo, Stephen J. Bush, Mark Fife, Paul M. Hocking, Georgios Banos, David A. Hume, Jim Kaufman, Richard A. Bailey, Santiago Avendano, Kellie A. Watson, Pete Kaiser, Mark. P. Stevens

## Abstract

*Campylobacter* is the leading cause of bacterial foodborne gastroenteritis in many countries. Source attribution studies unequivocally identify the handling or consumption of contaminated poultry meat as the primary risk factor. One potential strategy to control *Campylobacter* is to select poultry with increased resistance to colonisation. We conducted genomic and transcriptomic analyses of commercial pedigree broilers exposed to *Campylobacter* to examine persistent colonisation of the caecum as a quantitative trait. 3,000 broilers were genotyped using a 50K single nucleotide polymorphism (SNP) array and imputed to 600K SNPs. Genotypes were analysed for associations with the number of viable *Campylobacter* in the caeca. Heritability of the trait was modest but significantly greater than zero (h^2^=0.11 ± 0.03). Genome-wide association analyses confirmed quantitative trait loci (QTL) on chromosomes 14 and 16 previously identified using the progeny of crosses of inbred lines differing in resistance, and detected two additional genome-wide significant QTLs on chromosomes 19 and 26. RNA-Seq analysis of the transcriptome of caecal tonsils from birds at the low and high extremes of *C. jejuni* colonisation phenotype identified differentially transcribed genes, mainly located within the QTL on chromosome 16 and proximal to the major histocompatibility complex (MHC) locus. We also identified strong *cis*-QTLs located within the MHC suggesting the presence of *cis*-acting variation in both MHC class I, class II and BG genes. Multiple other *cis-*acting variants were identified in association with key immune genes (*COPS3, CCL4, CR1L, C4BP, PLGR*) in the other QTLs. Pathway and network analysis implicated cooperative functional pathways and networks in colonisation, including those related to antigen presentation, innate and adaptive immune responses, calcium, and renin-angiotensin signalling. While co-selection for enhanced resistance and other breeding goal traits is feasible, the frequency of resistance-associated alleles was high in the population studied and non-genetic factors significantly influence *Campylobacter* colonisation in poultry.

**Author summary:** *Campylobacter* infection is estimated to cause 95 million illnesses in people worldwide each year. Human infections mostly involve gastroenteritis, but can have severe complications. The handling or consumption of contaminated poultry meat is a key risk factor for human campylobacteriosis. The bacteria reach high numbers in the intestines of chickens reared for meat (broilers) and are frequently found on carcasses after slaughter. Effective vaccines against *Campylobacter* are not yet available, and treatments to reduce carcass contamination (e.g. chlorination) are not acceptable in some markets. One alternative is to breed for chickens with improved resistance to *Campylobacter* colonisation. To test the feasibility of this option in commercial birds, we analysed the genetic make-up of 3,000 pedigree broilers and determined the number of *Campylobacter* in their gut. There were associations between specific regions of the chicken genome and resistance to *Campylobacter*. Within some of these regions, expression of certain genes differed between birds at the low and high extremes of *Campylobacter* colonisation, providing a potential explanation for genetic variation in resistance. Selection of poultry with increased resistance to *Campylobacter* colonisation may be a complementary strategy to improved biosecurity, management, handling and processing procedures to reduce the burden of *Campylobacter* on human health.

## Introduction

Human campylobacteriosis exerts profound societal and economic costs. The World Health Organisation estimated that *Campylobacter* caused 95 million illnesses, 21,000 deaths and loss of 2.1 million disability-adjusted life years globally in 2010 [1]. In the United Kingdom alone, there were 63,946 laboratory-confirmed human cases in 2017, most of which were due to *C. jejuni* The ratio of unreported cases of human campylobacteriosis to those captured by national surveillance in the United Kingdom is estimated to be 9.3 to 1 [2] therefore the true burden may exceed half a million cases per annum at a projected median cost to the national health service of £50 million [3]. Human infections typically involve acute gastroenteritis, however debilitating sequelae may occur including Guillain-Barré syndrome (GBS) and other inflammatory neuropathies [4]. Poultry are the main reservoir of human campylobacteriosis and up to 80% of human cases may be attributable to the avian reservoir as a whole [5, 6]. During 2016-2017, a United Kingdom-wide survey indicated that 54% of fresh retail chicken was contaminated with viable *Campylobacter* [7]. The number of *C. jejuni* in the caeca of chickens frequently exceeds 10^8^ colony-forming units/gram and carcass contamination at slaughter can be difficult to avoid [8, 9]. Quantitative risk assessments predict that even a relatively modest 2 log_10_ reduction in the number of *Campylobacter* on broiler carcasses could reduce the incidence of human disease due to contaminated chicken by up to 30-fold [10]. A pressing need therefore exists for strategies to reduce the entry of *Campylobacter*-contaminated poultry meat into the food chain.

As effective vaccines and treatments for pre-slaughter control of *Campylobacter* in poultry are lacking, much interest exists in the potential for breeding chickens with improved resistance to intestinal colonisation by *C. jejuni*. The scope for genetic control of *Campylobacter* in poultry has been demonstrated in commercial broiler lines that vary in resistance [11-17] and inbred layer lines exhibiting heritable differences in *C. jejuni* colonisation following experimental inoculation [18, 19]. Breeding for *Campylobacter* resistance may also benefit avian intestinal health and productivity. In some commercial broiler chicken lines, experimental inoculation of *C. jejuni* elicits damage to the intestinal mucosa, diarrhoea and impaired weight gain [20, 21].In some birds, *Campylobacter* colonisation may impact intestinal barrier function [22], nutrient absorption and transporter expression [21, 23], the translocation of *Escherichia coli* to extra-intestinal organs [24], and may lead to a dysbacteriosis. In other studies, where *Campylobacter* inoculation was via natural exposure mimicking field exposure, there was no association between *Campylobacter* levels and bird growth rate [25] or gut pathology [26], and selection for *Campylobacter* resistance remained compatible with other breeding goals [26].

A previously published genome-wide association study (GWAS) on *C. jejuni* intestinal colonisation, where phenotypes were analysed as a binary trait in a novel dual-purpose chicken breed, revealed a resistance-associated locus on chromosome 11 near the *CDH13* gene encoding T-cadherin, and a second candidate locus on chromosome 5 was identified close to calmodulin (*CALM1*), a calcium-activated modulator of cadherin function [17]. Both genes differed in relative expression in a manner associated with resistance [17]. Studies in inbred layer lines 6_1_ and N found heritable differences in caecal colonisation by various *C. jejuni* strains [18]. Initial low-resolution mapping studies using reciprocal (6_1_♀×N♂) and (N♀×6_1_♂) F1 crosses and the progeny of an (N♀x6_1_♂) F1♂ x N♀ backcross indicated that resistance was controlled by a single autosomal dominant locus [18], but subsequent analysis of a backcross population using 1,243 fully-informative single nucleotide polymorphism (SNP) markers revealed quantitative trait loci (QTL) on chromosomes 7, 11 and 14 [19]. Using a ninth-generation advanced intercross (6_1_xN) line and a 600K genome-wide SNP array, the location of the QTL on chromosome 14 was confirmed and refined, and additional QTLs were identified on chromosomes 4 and 16, indicating potential involvement of the Major Histocompatibility Complex (MHC) region [19]. QTL for resistance of chickens to enteric carriage of *Salmonella* have been reported at the same regions of chromosome 14 [27] and 16 [27-29] that were implicated in host resistance to *C. Jejuni* colonisation. Analysis of caecal gene expression in chicken lines of varying susceptibility to *Campylobacter* colonisation has identified transcriptional signatures associated with differential resistance, including genes influencing the immune response [13-16].

In the present study, we conducted a comprehensive genome-wide association study on a commercial pedigree broiler population exposed to *Campylobacter* naturally present in the litter. The genomic architecture of resistance was analysed using imputed high-density SNP genotyping, and resistance was further characterised by transcriptome analyses of intestinal tissue from birds of the predicted resistant or susceptible genotypes at the corresponding extremes of caecal *Campylobacter* colonisation phenotype. Our data provide valuable insights into the prospects for genetic control of *Campylobacter* in poultry.

## Results

### Descriptive statistics and genetic parameters affecting *Campylobacter* levels

The mean number of caecal *C. jejuni* detected ± standard deviation was 7.057 ± 1.023 log_10_ colony-forming units (CFU) per gram (g), with a maximum of 10.64 log_10_ CFU/g and minimum of 2 log_10_ CFU/g, equal to the limit of detection by direct plating. Sex had a marginal effect on *Campylobacter* levels (*P* <0.05), with males having higher *Campylobacter* load (7.178 ± 0.034 log_10_ CFU/g) compared to females (6.930 ± 0.032 log_10_ CFU/g). *Campylobacter* levels showed seasonal variability, with date of sampling having a significant effect (*P* <0.05), while body weight did not have a significant effect on the trait. No significant maternal effects were identified. Modest, but statistically significant heritability for caecal *Campylobacter* level was estimated (h^2^ = 0.11 ± 0.03).

### Genetic association analyses

From 50K SNP genotype data obtained for 3,000 broilers, genotypes were imputed to 600K SNP for 2,718 birds. 282 samples failed the pedigree integrity testing prior to imputation and were removed. Imputation was not possible on chromosome 16 due to the complexity of the MHC region. These data were analysed using two genome-wide association methodologies: a genome-wide association study (GWAS) for single SNPs and regional heritability mapping (RHM) to consider associations with genomic regions comprising multiple consecutive SNPs. Using the SNP data on chromosome 16 the MHC haplotypes of these birds were assembled and a haplotype analysis was performed.

#### Genome-wide association study

Manhattan and quantile-quantile (Q-Q) plots for the GWAS results using the 50K and the imputed 600K arrays are shown in Fig. 1a and1b, respectively. SNPs significantly associated with log-transformed caecal *Campylobacter* levels, after a Bonferroni correction for multiple testing, are shown in Table 1. Multi-dimensional scaling analysis revealed no population substructure in these commercial broilers. GWAS analysis using the 50K SNP DNA array identified one SNP on chromosome 16 significantly associated with the log-transformed number of *Campylobacter* in the caeca at the genome-wide level and another ca. 100 SNPs on the same chromosome with a suggestive genome-wide significant association with the trait. All the significant SNPs were in high linkage disequilibrium (LD) and located within the same LD block, in the same MHC region (S1. Fig). Three MHC haplotypes were constructed. The recombination events were limited with only one event identified in the *TRIM* region of MHC of one sample. One of the MHC haplotypes (AA) was detected at much higher frequency (88%), compared to the other two (BB = 0.5% and BA = 11.5%). The ensuing MHC haplotype analysis identified statistical significant (*P*<0.05) associations between the MHC haplotypes and the log-transformed number of *Campylobacter* in the caeca. The prevalent MHC haplotype AA was associated with colonisation resistance accounting for 1 log_10_ difference in the *Campylobacter* levels compared to the susceptible BB haplotype; a significant favourable dominance deviation was also observed for the heterozygous haplotype AB (Fig. 2).

**Table 1.** List of SNPs significantly associated with log-transformed caecal *Campylobacter* levels in commercial chickens.

| SNP array | SNP marker | Location (Chr(bp)) | GWAS (P-value) | Additive effects (P-values) | Dominance effects (P-values) | Phenotypic Variance explained by the SNP locus | p | q |
| --- | --- | --- | --- | --- | --- | --- | --- | --- |
| 50K | **AX-76346701** | 26 (4644002) | 1.15E-06 | -0.19(0.10) | **0.46(0.05)** | 0.05 | 0.88 | 0.12 |
| 50K | **AX-75852372\*** | 16(70922) | 1.16E-06 | **0.65(0.001)** | -0.37(0.07) | 0.06 | 0.94 | 0.06 |
| 50K | AX-75852319 | 16(65899) | 1.17E-06 | **0.62(0.02)** | -0.34(0.09) | 0.06 | 0.94 | 0.06 |
| 50K | AX-75778268 | 14(12417411) | 1.18E-05 | **0.38(0.038)** | -0.30(0.07) | 0.03 | 0.93 | 0.07 |
| 50K | AX-76339934 | 26(3224490) | 1.50E-05 | -0.12(0.20) | **0.40(0.05)** | 0.03 | 0.89 | 0.11 |
| 50K | AX-75798135 | 14(4937371) | 7.09E-06 | 0.07(0.30) | 0.15(0.20) | 0 | 0.93 | 0.07 |
| 600K | AX-76346702 | 26(4644456) | 4.47E-07 | 0.13(0.24) | **-0.38(0.05)** | 0.04 | 0.88 | 0.12 |
| 600K | AX-75798142 | 14(4939239) | 1.32E-06 | 0.05(0.39) | 0.09(0.36) | 0 | 0.93 | 0.07 |

SNP markers highlighted in bold were significant at the genome-wide level after Bonferroni correction.
\*More than 100 SNP markers on chromosome 16 were significant at the suggestive significant genome-wide level.

**Fig 1.**
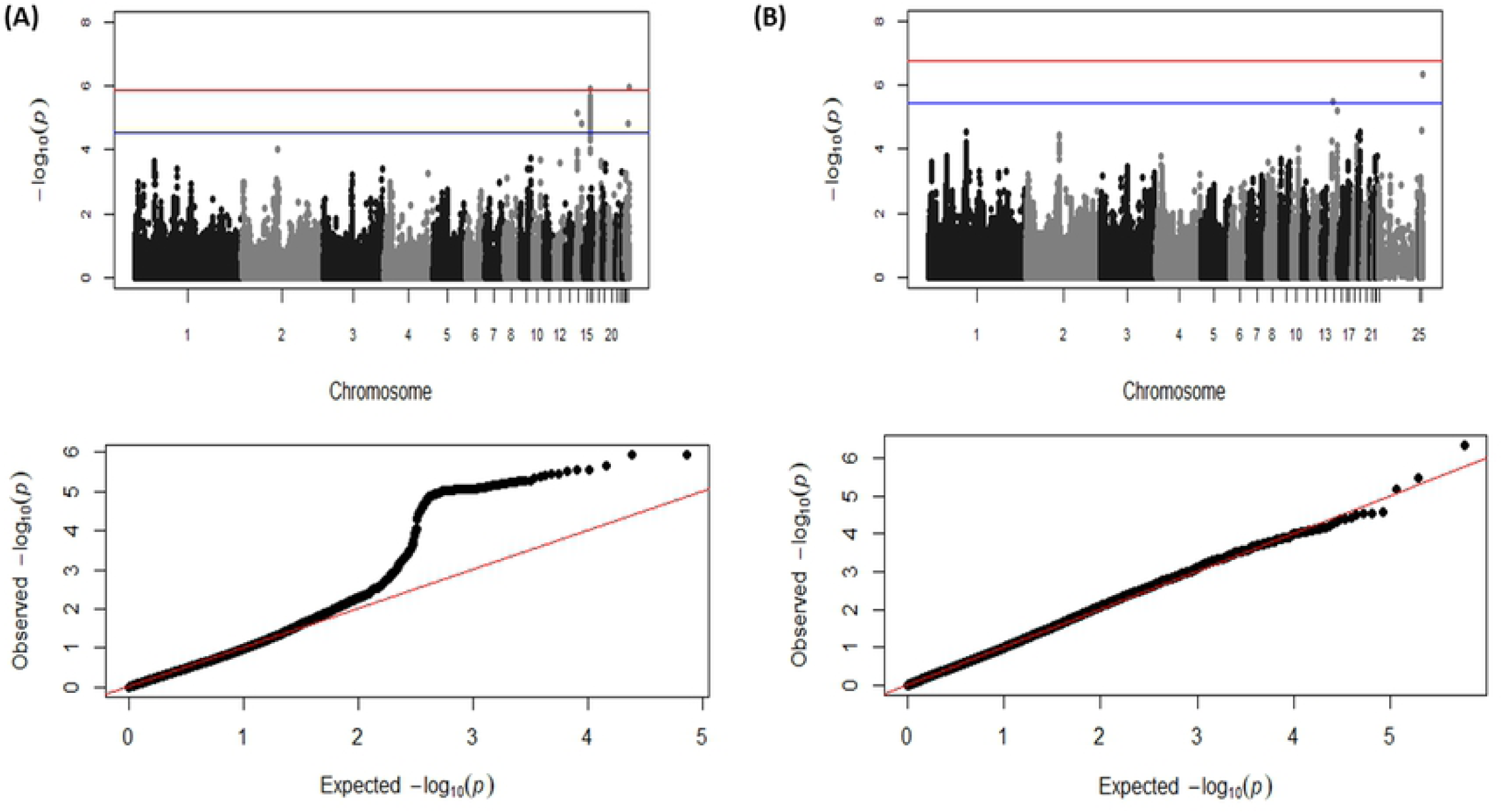
Manhattan plots and Q-Q plots displaying the GWAS results for chicken *Campylobacter* colonisation resistance using the 50K (A) and the imputed 600K (B) HD arrays. (i) Genomic location is plotted against – log_10_(P) in the Manhattan plot. Genome-wide (*P* < 0.05) and suggestive genome-wide thresholds are shown as red and blue lines, respectively. (ii) Q–Q plot of observed *P* values against the expected *P* values for *Campylobacter* caeca load (log-transformed CFU of *Campylobacter* per gram of caeca content).

**Fig 2.**
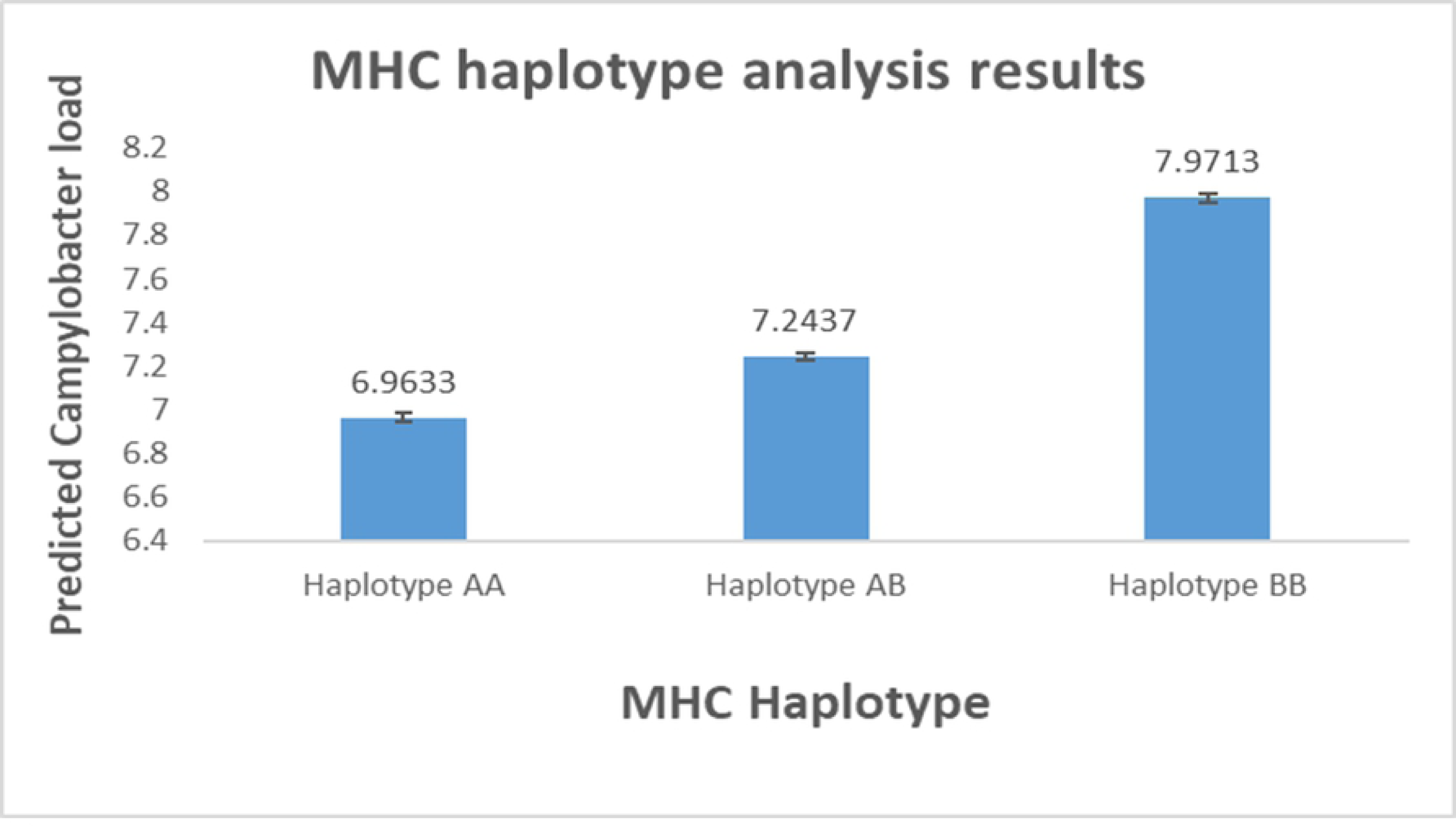
MHC haplotype analysis results. *Campylobacter* caecal load (log-transformed CFU of *Campylobacter* per gram of caeca content) is plotted against the MHC haplotypes identified in the commercial chickens.

Additionally, one SNP with a genome-wide significant association and another one, located 1.4Mb away, with a suggestive association were identified on chromosome 26. Similarly, two SNPs identified on chromosome 14, albeit 7.5Mb apart from each other, crossed the suggestive genome-wide significant threshold. The significant SNPs on chromosomes 14 and 26 were not in LD. The GWAS analysis using the imputed 600K SNP array information confirmed the associations on chromosomes 14 and 26.

The additive and dominance genetic effects, and the proportion of the total phenotypic variance explained by each of these SNPs are summarised in Table 1. The SNPs on chromosome 14 and 16 had a significant additive effect (ranging from 0.3 to 0.6 log_10_ CFU/g) while the SNPs on chromosome 26 had a significant dominant effect (ranging from 0.3 to 0.4 log_10_ CFU/g) on *Campylobacter* levels. The significant SNPs in the QTL region on chromosome 16 accounted for 6% of the phenotypic variance, while collectively all the SNPs in the three candidate regions accounted for 17% of the phenotypic variance of caecal *Campylobacter* levels.

#### Regional heritability mapping

A Manhattan plot and Q-Q plot for the RHM analysis are shown in S2 Fig. Details of the significant SNP windows are presented in S1 Table. RHM confirmed the significant associations on chromosome 16 previously identified by the GWAS. Moreover, RHM detected one more suggestive significant association on chromosome 19.

### SNP and candidate region annotation

Most of the significant SNPs identified in the GWAS analysis were located upstream (40 %) or downstream (34 %) of predicted genes or within introns (19 %). However, eight of the SNPs on chromosome 16, one SNP on chromosome 26 and one SNP located within the significant RHM SNP-window on chromosome 19 were found within exons. The exonic variants on chromosome 16 were located within *TRIM10* (tripartite motif 10), *TAP1* (transporter associated with antigen processing 1), *RACK1/GNB2L1* (receptor for activated C kinase 1), *TRIM27* (tripartite motif 27), and *TRIM32* (tripartite motif 32). The SNP on chromosome 19 corresponds to a synonymous variant within *HIP1* (huntingtin interacting protein 1) and the SNP on chromosome 26 corresponds to a missense variance within *ADORA3* (adenosine receptor 3).

The candidate QTL regions for caecal *Campylobacter* levels contained a relatively small number of genes, collectively comprising 173 annotated protein-coding genes, 7 microRNAs and 2 snoRNAs. Details of the genes and non-coding RNAs located in the candidate regions are presented in S2 Table.

### Transcriptomic analyses

As many traits are associated with altered expression of genes within QTLs [30], we performed RNA-Seq analysis of the caecal tonsil transcriptome of 23 broilers to identify expression QTLs (eQTLs) and potential allelic imbalance of candidate genes within the regions associated with *Campylobacter* resistance identified with GWAS and RHM. The birds used were selected to represent combinations of predicted resistant or susceptible genotypes (based on the identified QTLs) that were correspondingly at the extremes (resistant n=9, susceptible n=7) as well as the average (n=7) of measured caecal *Campylobacter* load (S3 Table).

#### Differential gene expression analysis

After false discovery rate (FDR, *P<*0.05) correction for multiple testing, 3 protein-coding genes were found to exhibit significant differential expression (Table 2). The three differentially expressed (DE) genes were located within the QTL region on chromosome 16 (*BF2, ENSGALG00000032221, ENSGALG00000024357*). The *BF2* gene is an MHC class I gene while the other two are both *BG*-like genes belonging to the butyrophilin family. In order to identify more subtle patterns of differential expression, a relaxed significance threshold of unadjusted *P* value of 0.001 was implemented and a total of 33 genes exhibited differential expression between high-, average-, and low-colonised birds at this threshold (Table 2, Fig. 3). Among these DE genes were several related with the immune response (*ILF2, ATG7, BG1, BF2, BF1, TAP1, ZNF692*). Interestingly, there were three DE *BF2* transcripts, two of which were downregulated (ENSGALT00000079478, ENSGALT00000077683) and the other (ENSGALT00000087837) upregulated in the resistant birds. There were also two DE *BF1* transcripts, both of which were upregulated in the resistant birds.

**Table 2.**
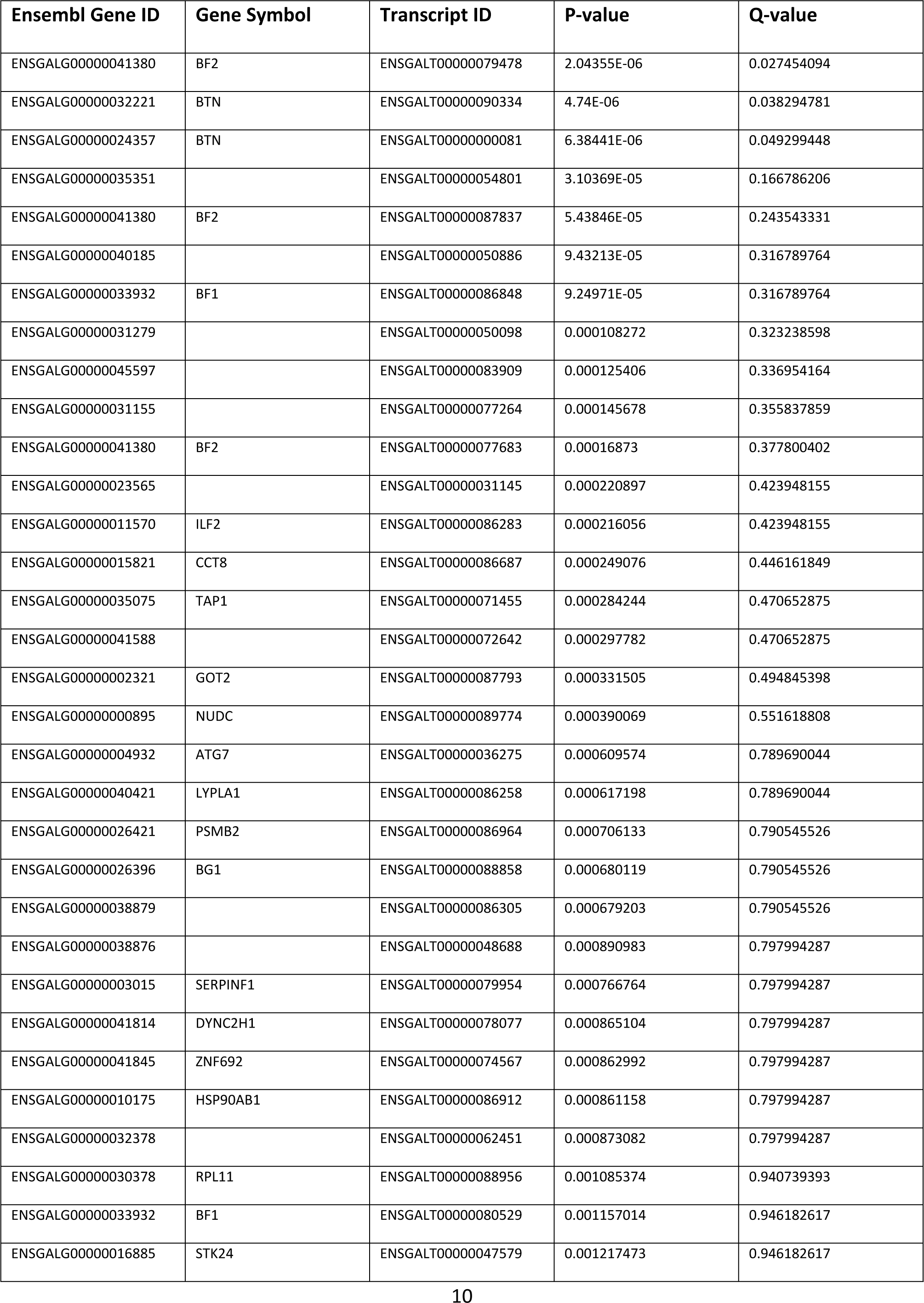
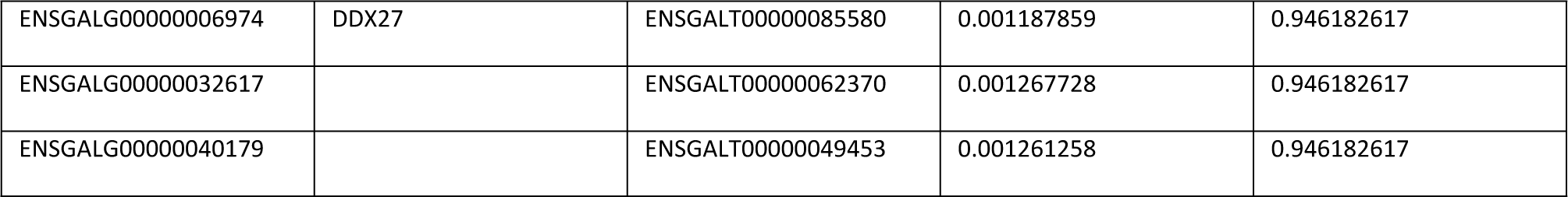
List of differentially expressed transcripts in the caecal tonsils of low, average and high *Campylobacter* colonised commercial chickens.

**Fig 3.**
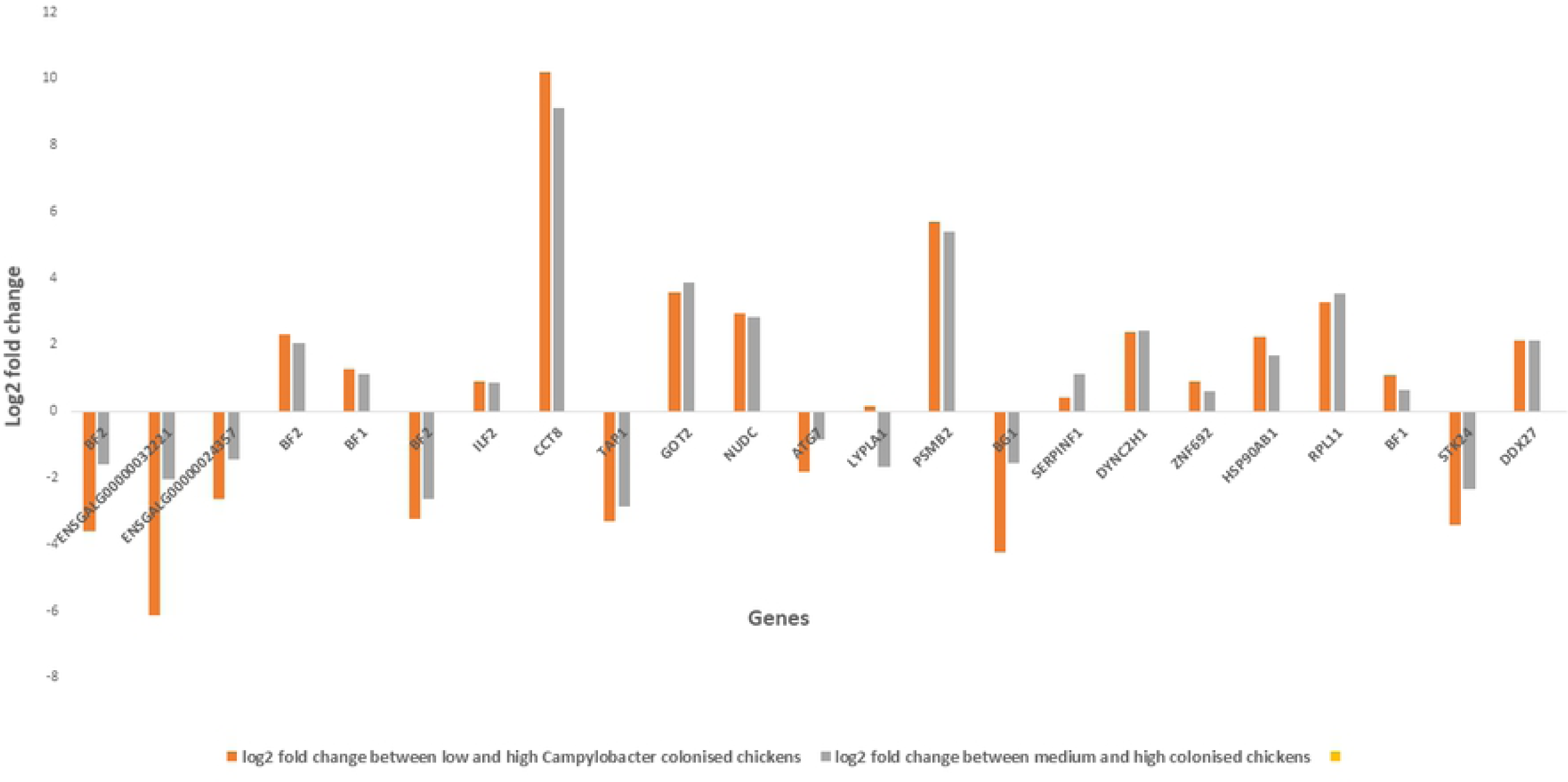
Differential expression analysis results. Differential expression of genes in chickens with different (low, average and high) *Campylobacter* colonisation levels. Each column represents relative gene expression levels in caecal tonsils of chickens. Expression level is shown as log2 fold change in expression of low and average colonised chickens relative to expression of high colonised chickens.

We performed separate *cis-* and *trans-* based eQTL analyses for the significant and suggestive significant SNP markers identified by the GWAS and RHM:

#### Cis-analysis

After false discovery rate (FDR, *P*<0.05) correction for multiple testing, we detected 102 significant *cis*-eQTL (S4 Table). Of those, 90 were associations between SNPs in high LD, located in the same QTL region on chromosome 16, and the expression of a single gene, BG-like antigen 1 (*BG1*) (Fig. 4A). This eQTL had a log_10_ allelic-fold-change of 2.03. Four more *cis*-eQTLs were identified for *ENSGALG00000032221* and three novel gene transcripts on chromosome 16, *ENSGALT00000065054, ENSGALT00000049453* and *ENSGALT00000085167*. Another three significant *cis*-eQTLs were detected within *TMEM11* (transmembrane protein 11) and the *COPS3* (COP9 signalosome subunit 3), two genes located within the QTL region on chromosome 14 (position 4,552,835-4,560,698 and 4,767,396-4,781,731, respectively) (Fig. 4B and 4C, S4 Table).

**Fig 4.**
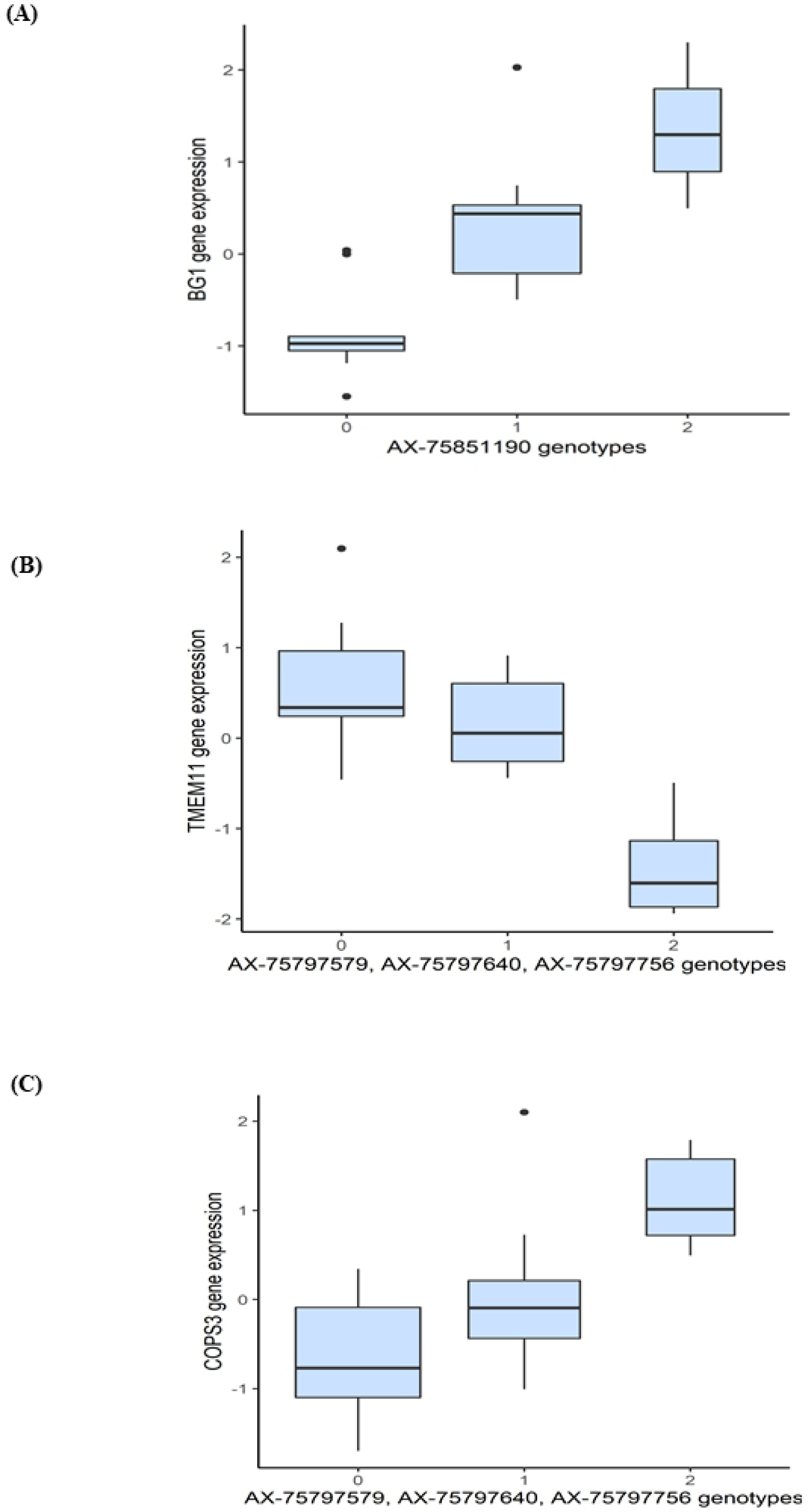
Expression QTL analysis results. Boxplots showing the expression of *BG1* (4A), *TMEM11* (4B) and *COPS3* (4C) genes depending on the genotypes of SNPs acting as cis-elements. On the x-axis, “0”, “1” and “2” represent the number of copies of the non-reference allele, and on the y-axis the expression of each gene (Transcripts Per Million, TPM), is represented after mean-centering and scaling.

#### Trans-analysis

We detected a total of 13 significant *trans*-eQTLs within the QTL region on chromosome 19 and two on chromosome 26 (S5 Table). Most of these predicted *trans*-acting elements are for genes related with metabolic processes. The *trans*-acting SNP on chromosome 26 is for a microRNA (gga-mir-1553) located on chromosome 7, close to the peak of a previously identified QTL for *C. jejuni* resistance identified using a back-cross population of inbred lines 6_1_ and N [19].

#### Allele-specific expression analysis

If an individual is heterozygous for a *cis*-acting SNP it is expected that the two alleles of the gene will be expressed unequally causing allelic expression imbalance. To verify the *cis*-QTLs detected above, and identify additional ones, we identified genetic variation within the QTL regions identified by the GWAS and RHM using the RNA-Seq data and performed allele-specific expression (ASE) analysis for all the SNPs located within these regions. Several significant ASE events were identified in all QTL regions (mean *P* value ≤ 0.05 with at least 4 heterozygous animals). Specifically, 14 significant ASE events were identified for 3 genes located on chromosome 14, 30 for 6 genes on chromosome 16, 11 for 2 genes and one microRNA (gga-mir-142) on chromosome 19, and 35 for 5 genes on chromosome 26 (S6 Table). A highly significant ASE event was identified on chromosome 14 (for the QTL located at 12MB) for chloride voltage-gated ion channel 7 (*CLCN7*). ASE results on chromosome 16 were consistent with the presence of a *cis*-acting polymorphism in *BG1*, with 6 SNPs in high-LD showing allelic imbalance (*P* < 10^−13^). Moreover, ASE analysis highlighted potential *cis*-acting polymorphisms for other genes of interest in the region, namely MHC class 1 (*BF1 and BF2*) and class 2 (*BLB2* and *BLB1*) (Fig. 5A-5C). Within *BF2*, 13 different SNPs showed highly significant ASE, with *P* values < 10^−305^; Fig. 5A). Chromosomes 19 and 26 also contain several immune-related genes showing significant ASE: angiopoietin-related protein 2 (*ANGPTL2*), C-C motif chemokine ligand 4 (*CCL4*), complement C3b/C4b receptor 1-like (*CR1L*), C4b-binding protein (*C4BP*), polymeric immunoglobulin receptor (*PLGR*) and BCL2 antagonist/killer 1 (*BAK2*).

**Fig 5.**
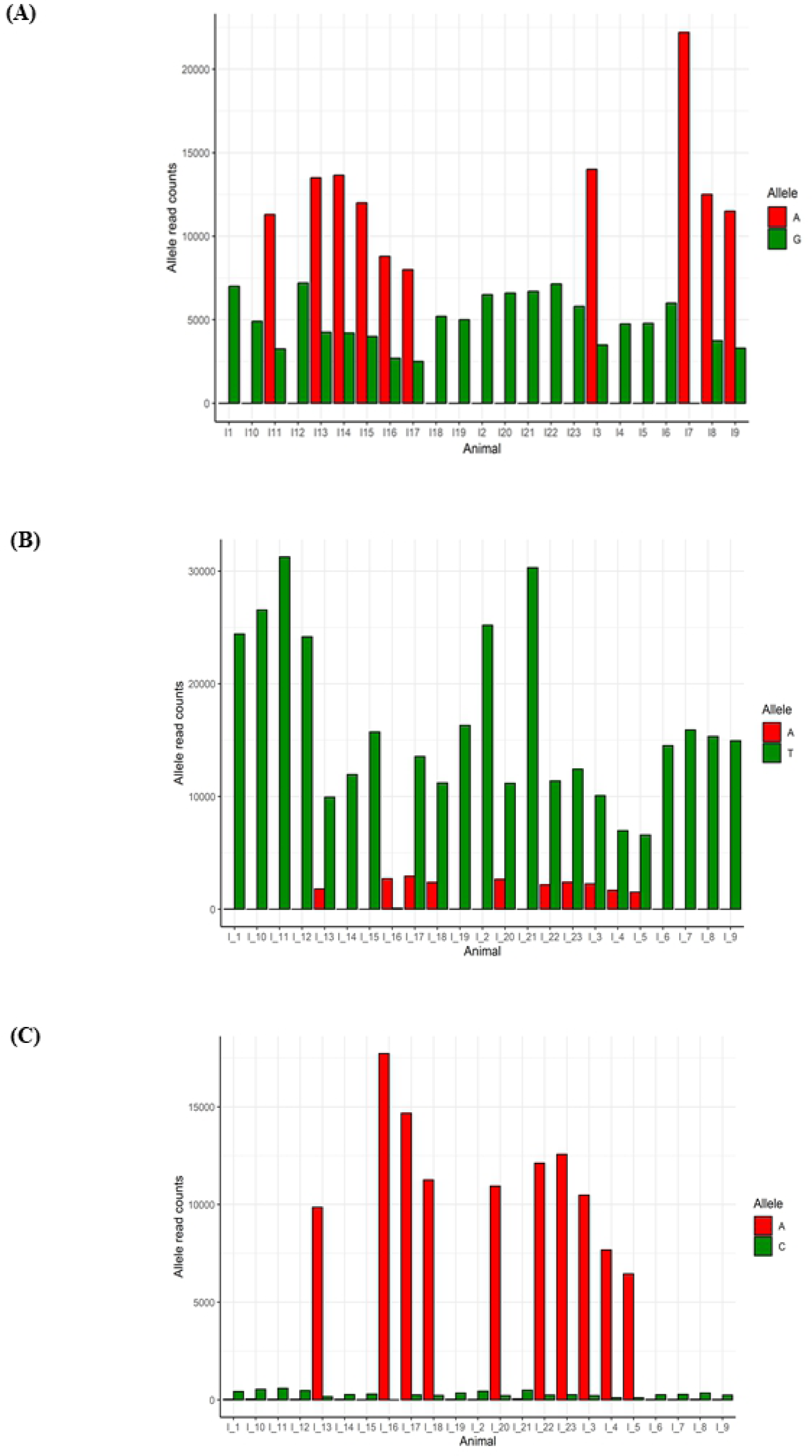
Allele specific expression (ASE) analysis results. Bar plots showing the allele specific expression results for *BF2* (5A), *BLB1* (5B) and *BLB2* (5C) genes. Each column represents gene expression levels, measured as read counts, for each allele (reference (red) vs non-reference (green) allele). Gene expression levels have been plotted against each individual animal.

#### Validation of selected differentially transcribed genes

Specific qRT-PCR assays were devised to validate the transcript levels measured by RNA-Seq using the same RNA samples. Four genes located in the MHC region on chromosome 16 (*BF2, BF1, ENSGALG00000032221* and *ENSGALG00000024357*), were found to be differentially transcribed in the caecal tonsils of birds of predicted resistant or susceptible genotypes with divergent caecal *Campylobacter* load, after adjusting for sex and seasonality. This confirmed that the expression of each gene differed significantly between resistant and susceptible birds (*P* ≤ 0.05 after Tukey’s HSD post-hoc test adjustment).

### Pathway, network and functional enrichment analyses

#### Pathway analysis using encoded genes in the candidate regions for *Campylobacter* resistance

Based upon the significant heritability estimate and the large amount of genetic variance accounted for by the identified SNPs, we reasoned that the corresponding QTL regions may contain genes contributing to common pathways associated with resistance to *Campylobacter* colonisation. We therefore identified the sets of annotated genes lying within QTL regions and sought evidence of gene set enrichment. Ingenuity Pathway Analysis (IPA) found these genes to be enriched for pathways involved in innate and adaptive immune responses, antigen presentation, inflammatory responses, calcium signalling, epithelial cell signalling and interactions (Fig. 6). Moreover, three networks of molecular interactions related to ‘immunological diseases’, ‘cell death and survival’, and ‘molecular transport and protein trafficking’ were constructed using the list of genes in the candidate regions (Fig. 7). We subsequently extracted the gene ontology terms for each of these genes and performed functional annotation clustering analysis. The genes were organised into 41 clusters, each given an enrichment score (ES). The first (ES = 4) and the second (ES = 3.5) clusters were both enriched for genes functionally annotated as involved in ‘antigen processing and presentation via MHC class I and class II molecules’ (including *BF1, BF2, BLB1, BLB2, DMB1, DMB2, TAP1*, and *TAP2*) (S7 Table).

**Fig. 7.**
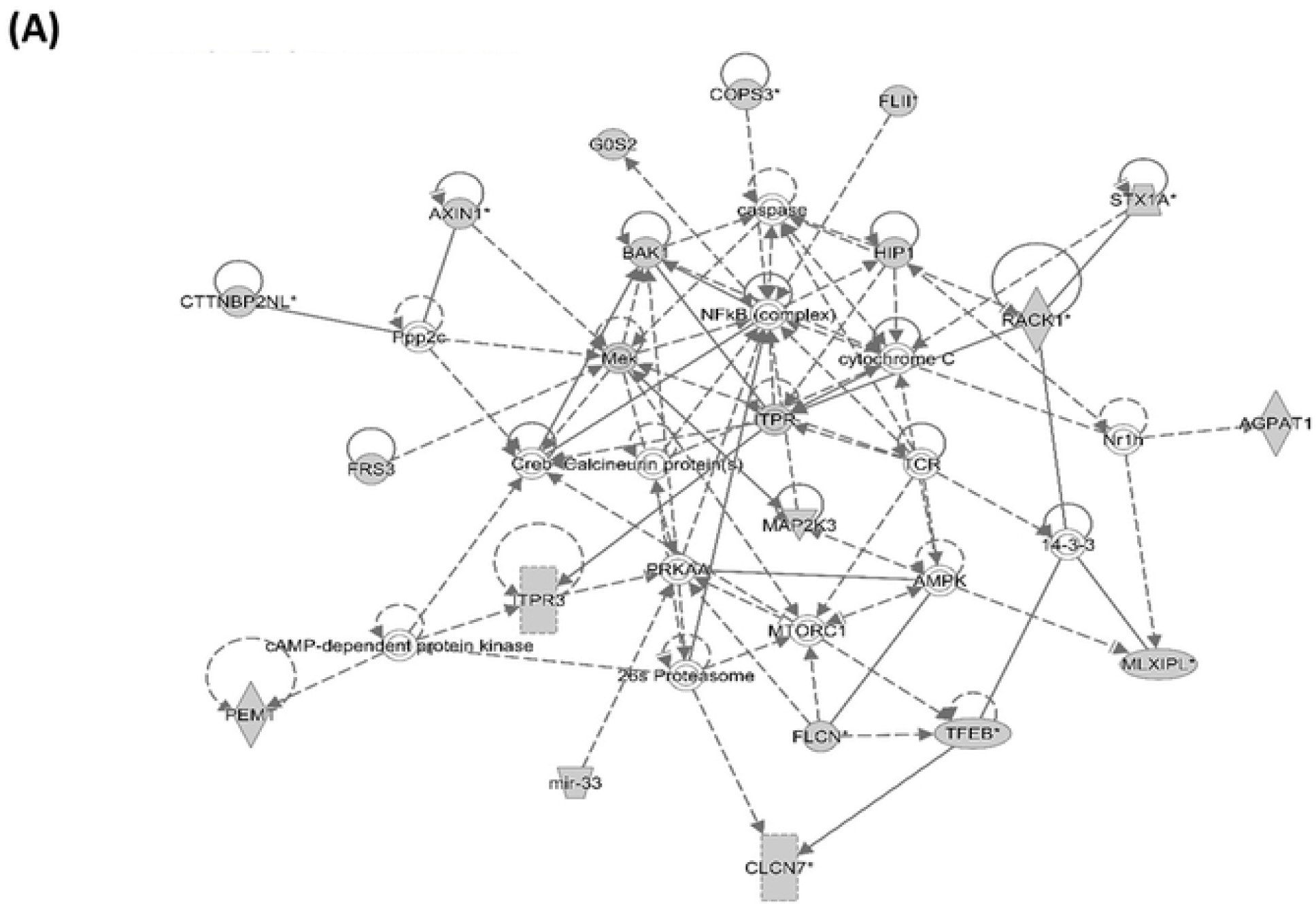
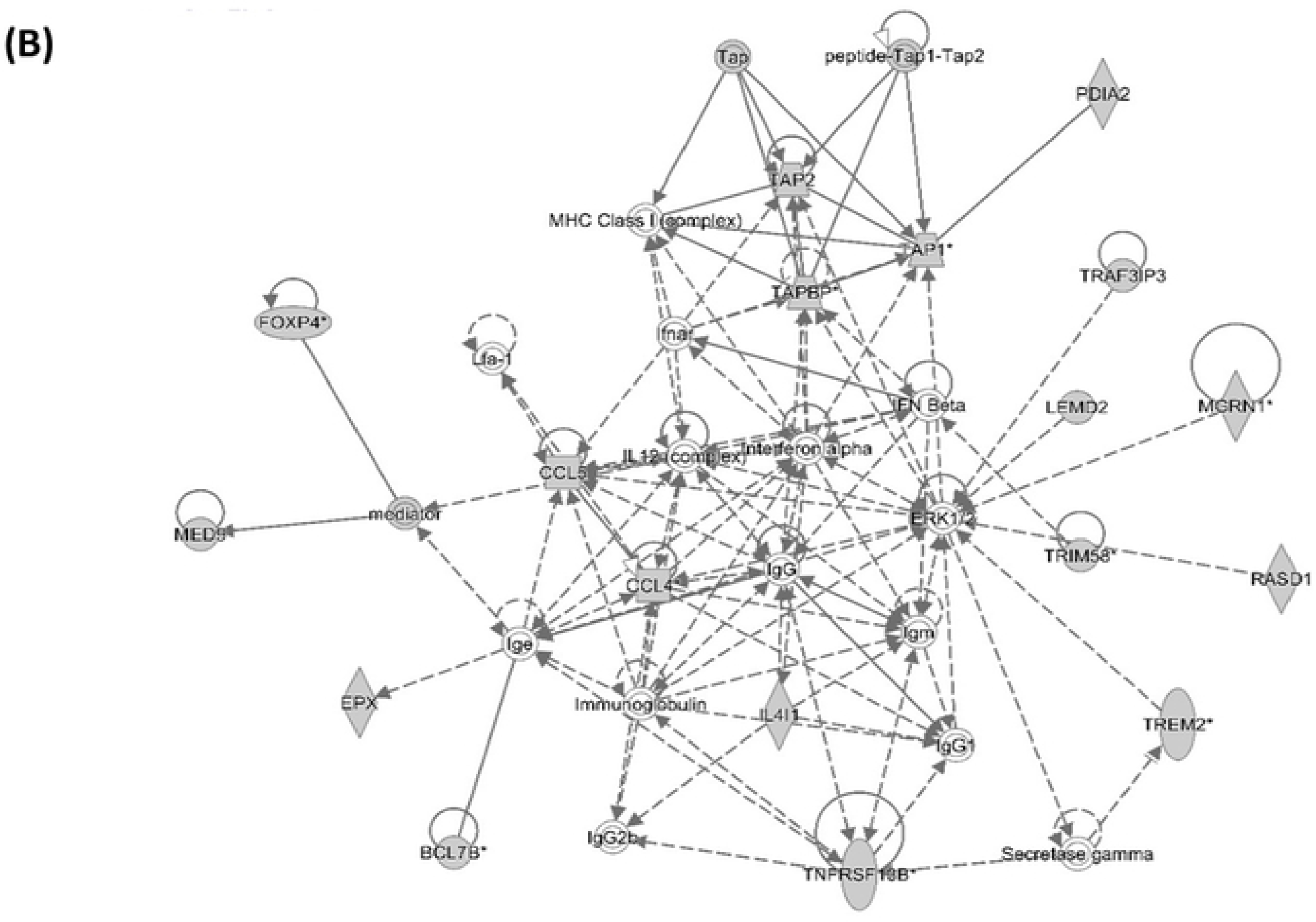
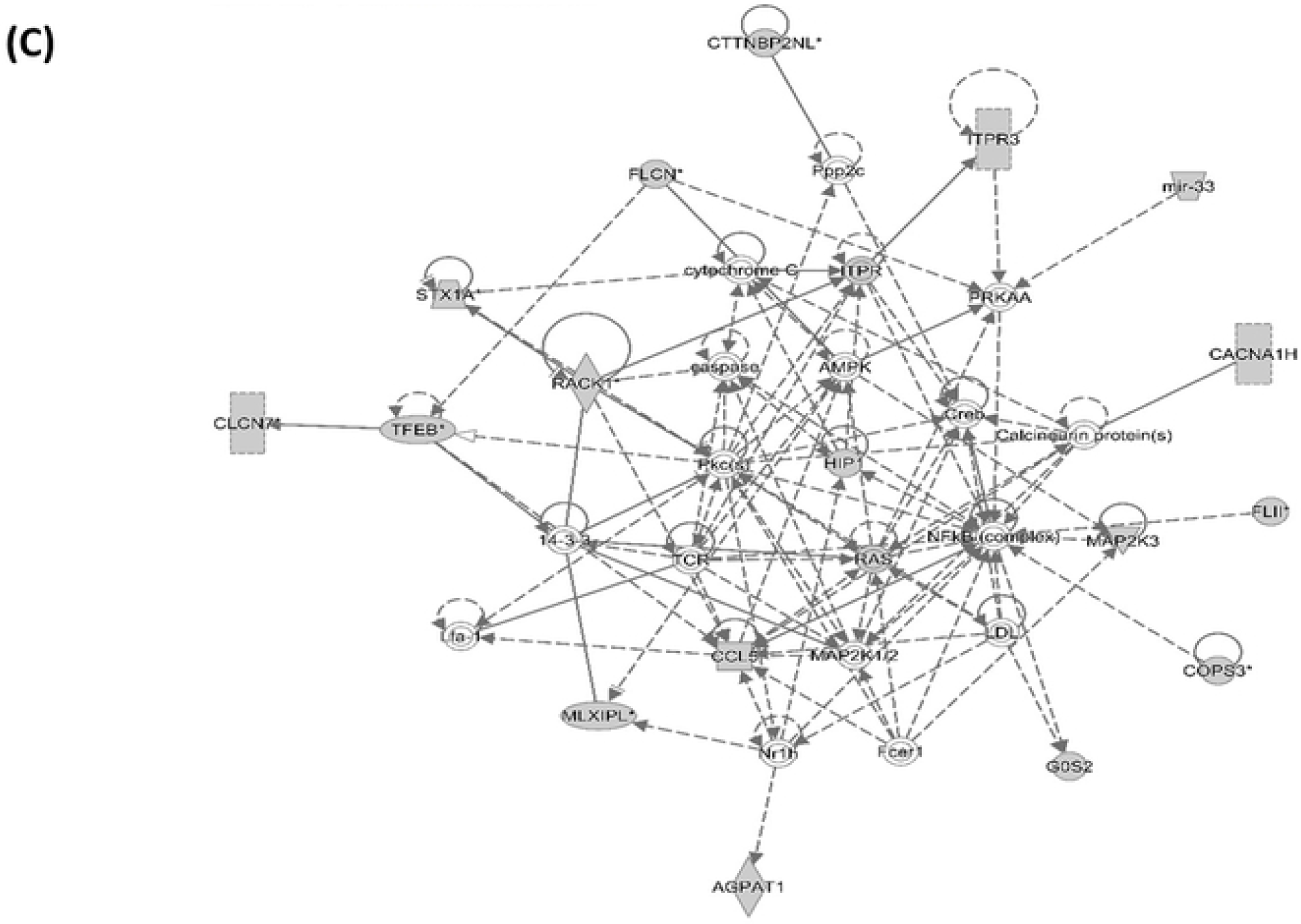
Network analysis using the IPA software. Three networks, related to immunological disease (A), cell death and survival (B), and molecular transport and protein trafficking (C) that illustrate the molecular interactions between the products of candidate genes selected from QTL regions for *Campylobacter* colonisation resistance in commercial chickens. Arrows with solid lines represent direct interactions and arrows with broken lines represent indirect interactions. Genes with white labels are those added to the IPA analysis because of their interaction with the target gene products.

#### Pathway analysis using DE genes in birds with different *Campylobacter* colonisation levels

Functional analysis of the DE genes using the IPA software showed significant enrichment for pathways related with immune response (interferon signalling, antigen presentation, immunodeficiency signalling) and metabolism (protein ubiquination, glutamate degradation) (S3 Fig.). Moreover, one network of molecular interactions related to “cell death and survival, and organismal injury and abnormalities” was constructed based on the DE genes (S4 Fig.). Functional annotation clustering of these genes uncovered significant enrichment (E.S. = 1.2) for one gene cluster related with immune response, defence response, response to stress, symbiosis, encompassing mutualism through symbiosis, interspecies interactions between organisms.

## Discussion

We sought to investigate the genetic basis of resistance of chickens to *Campylobacter* colonisation and evaluate the potential for selective breeding of poultry with enhanced resistance to control *Campylobacter* at farm level. Using samples from 3,000 commercial chickens exposed to *Campylobacter*, we detected heritable variation associated with caecal *Campylobacter* levels and identified genomic markers and regions associated with colonisation. Candidate genes, *cis*- and *trans*-acting elements, canonical pathways and networks, and MHC haplotypes that were implicated in resistance to *Campylobacter* colonisation were also identified.

We estimated significant heritability (h^2^=0.11) for caecal *Campylobacter* colonisation. This was lower compared to a previous estimate for this trait using the progeny of crosses of inbred White Leghorn chicken lines with differing resistance to *Campylobacter* colonisation (h^2^=0.25) [19]. The difference is likely attributable to the use of field data on naturally colonised broilers in the present study compared to experimental challenge of inbred lines. The heritability of resistance to *Campylobacter* colonisation is similar to that observed for other livestock pathogens and diseases, such as bovine tuberculosis (h^2^=0.09-0.17) [31, 32] and bovine and ovine mastitis (h^2^=0.10-0.20) [33], where the development of genetic evaluations to guide breeding decisions was deemed feasible.

In a previous study of the same commercial chicken population, *Campylobacter* colonisation levels were not significantly phenotypically and genetically correlated with key production traits such as body weight, nutrient absorption and gut health; this highlights that the presence of *Campylobacter* in the caeca of chickens was not detrimental to the birds studied and that co-selection for *Campylobacter* colonisation resistance with other breeding goals is feasible [26]. However, the low heritability estimates indicate that a large proportion of phenotypic variance in *Campylobacter* colonisation is determined by non-genetic factors that merit further investigation. Moreover, the high frequency of resistance-associated alleles in the studied population of commercial birds suggests limited scope for improvement, albeit commercial broiler lines from other breeding programmes may benefit from selection guided by the data presented here.

In the present study, we assumed a uniform exposure of birds to *Campylobacter* during the 16 months of sampling. A seasonal, batch and sex effect on *Campylobacter* colonisation was detected and fitted in the GWAS, eQTL and differential expression models of analysis to adjust for these sources of systematic variation. Season has been previously reported to affect the colonisation phenotype in chickens [34, 35], with this linked to an elevated incidence of human campylobacteriosis during summer [36]. The basis of this seasonal effect is not entirely clear [37]. Moreover, while *Campylobacter* was routinely detected in the environment of the birds sampled, we cannot preclude the possibility that the bacterial species and sequence types present varied over time.

Consistent with a previous report of paternal effects on caecal *C. jejuni* colonisation in broilers [12], we detected a significant effect of sex on the colonisation phenotype, with males having higher mean caecal counts of *Campylobacter*. Male susceptibility to *Campylobacter* has been also reported in human and mouse studies [38, 39]. Sex-related differences in immune response and survival rate of broiler chickens have been reported for a range of pathogens in chickens [40]. Male broilers were found to be more susceptible to infectious disease and this was attributed to a less efficient immune response compared to females [40]. Moreover, there are differences in gene expression and responsiveness to bacterial lipopolysaccharide between macrophages from males and females that have been attributed to the lack of dosage compensation of the genes on the Z chromosome, which includes the interferon cluster [41]. Apart from seasonality and sex, other non-genetic factors may explain the observed variation in *Campylobacter* colonisation, including strain variation [42], the time and level of exposure relative to sampling [43], coinfections [44], variation in the gut microbiota [45, 46], and diet and feed intake [47, 48]. Our results should be interpreted in the context of the limitations and advantages of field-based genome-wide association studies [49, 50]. Compared to controlled challenge experiments, unknown and uncontrolled exposure to non-genetic factors may reduce the power of a field study but do not constitute a fatal flaw in demonstrating host genetic differences in resistance [49]. Moreover, the demonstration of heritable resistance in field studies that simulate commercial practice is highly relevant to the production system into which selectively-bred birds would be introduced.

In line with our previous findings using the progeny of crosses of inbred chicken lines [19], the major histocompatibility complex region on chromosome 16 was implicated in resistance to *Campylobacter* colonisation in commercial broilers. Using genomic data we were able to identify a strong QTL in the MHC region explaining most of the trait-associated genetic variation, and the QTL overlapped with expression QTLs detected by RNA-Seq analysis of caecal tonsil tissue from birds at the extremes of the colonisation phenotype. Within this QTL region, 100 SNP markers were found in high LD and collectively corresponded to three MHC haplotypes. These haplotypes were relatively stable, since only one recombination event was identified in the *TRIM* region of MHC, and they were associated with distinct colonisation phenotypes, with the more prevalent one associated with colonisation resistance accounting for 1 log_10_ difference in the *Campylobacter* levels.

Despite the MHC region being polymorphic and repetitive, making it challenging to identify causative genes and mutations underlying disease resistance, our analyses revealed a number of candidate genes for *Campylobacter* resistance that warrant further investigation. Specifically, the eQTL and ASE analyses showed evidence for *cis-acting* elements related with the expression of the BG-like antigen 1 (*BG1*) gene, major (*BF2*) and minor (*BF1*) MHC class I genes, the major (*BLB2)* and minor (*BLB1*) MHC class II beta chain genes. In addition, the *BF2, BF1, BG1* and two BG-like genes were found to be differentially expressed in chickens with different caecal *Campylobacter* levels. The major MHC class II beta chain gene (*BLB2*) gene is widely expressed at high levels in hematopoietic cells, whereas the minor MHC class II beta chain gene (*BLB1*) is generally poorly expressed, although highly expressed in spleen, intestinal epithelial cells, and particularly the caecal [51]. The *CIITA* transactivator gene that controls expression of MHC class II genes [52] has been found to be differentially expressed in chickens with high and low *C. jejuni* colonisation levels in a previous experimental RNA-Seq study of the caecal tissue [16]. In that study, MHC class I genes *BF1* and *BF2* were also found to be differentially expressed in the caeca of chickens with high and low *C. jejuni* levels [16]. Furthermore, similar to our findings, MHC-related *BG* genes have also been reported to be differentially transcribed in the spleen of two chicken lines that differ in susceptibility to *C. jejuni* colonisation. The *BG* region of MHC is very repetitive, and it is therefore difficult to distinguish specific *BG* genes due to copy number variation [53].

In the present study, network analyses detected interferon signalling among the pathways associated with resistance. Interferon-γ (IFN-γ) has been reported to be induced following *Campylobacter* infection of avian cells [54] and chickens in a breed-dependent manner [55], and may underlie breed-specific differences in gut inflammation and pathology [20, 56]. Furthermore, multiple interferon-related genes were found to be differentially expressed in the caecal transcriptome of chickens with high and low *C. jejuni* colonisation levels in a previous RNA-Seq study [16]. Interestingly, the major class I and II molecules, as well as other MHC related genes have internal ribosome entry site (IRES) which respond to IFN-γ and therefore their differential expression may be subject to interferon regulation. Furthermore, our recent analysis of whole genome sequence of commercial broilers and layers, and the transcriptome of isolated macrophages from a broiler-layer F2 sibling backcross, also revealed substantial differences in the expression of interferon-regulatory factors (IRF) family members as well as in the *BLB1* and *BLB2* genes between individual birds [57] that could underlie this phenotype.

The present study identified two distinct QTLs on chromosome 14, both located within the interval of a previously identified QTL for *Campylobacter* resistance using a backcrossed ([6_1_ x N] x N) population of inbred lines 6_1_ and N [19]. One of these QTLs (located at 12 Mb) overlaps a QTL identified for the same trait using a ninth generation advanced intercross population of these lines [19], as well as a QTL for resistance to *Salmonella* colonisation in chickens [27], suggesting that a mechanism of resistance common to both pathogens may exist. CREB binding protein (*CREBBP*), a key immune regulatory protein implicated in *Salmonella* resistance in chickens [58], lies in close proximity to the marker of this QTL. In the present study, a mutation (14:12556836, C to T, splice donor variant) with a predicted high impact on the encoded protein of this gene was significantly (*P*<0.05) associated with *Campylobacter* colonisation resistance (data not shown). Furthermore, pathway analysis in the present study confirmed the enrichment for *CREB* signalling reported previously in inbred lines [19]. The other QTL region (located close to 5Mb) on chromosome 14 in the present study overlapped with an expression QTL for *Campylobacter* resistance. Specifically, the SNP marker significantly associated with *Campylobacter* resistance was also a *cis-*acting element for two genes (upregulates *COPS3* and downregulates *TMEM11*). The protein encoded by COP9 signalosome complex subunit 3 gene possesses kinase activity that acts as a site for complex phosphorylation of many regulators involved in signal transduction such as I-kappa-B-alpha, p105, and c-Jun [59]. This protein is part of a complex that plays a key role in diverse cellular processes including cytokine signalling and antigen induced responses [60].

Several immune-related genes in the QTL regions on chromosomes 19 and 26 showed evidence of allele-specific expression. Among these genes were the polymeric immunoglobulin receptor (*PLGR*) which is highly expressed in intestinal epithelial cells and mucosa, and plays a crucial role in the transcytosis of polymeric soluble immunoglobulins and immune complexes to the gut mucosal surface [61]. Genes and pathways involved in the immunoglobulin production and function were reported to be upregulated in chickens relative resistance to *C. jejuni* [16]. *PLGR* has been associated with intestinal immune defence against the lumen-dwelling parasite *Giardia* in mice [62]. The QTL for *Campylobacter* resistance identified on chromosome 26 encompasses the calcium/calmodulin-dependent protein kinase IG which belongs to a calcium-triggered signalling cascade and phosphorylates the transcription factor CREB (https://www.uniprot.org/uniprot/Q96NX5). A previous GWAS study of *C. jejuni* resistance identified a suggestive significant association proximal to the calmodulin gene [17]. Intracellular calcium levels in the intestinal epithelium are affected by *C. jejuni* in some lines [23], however the extent to which this affects bacterial colonisation, or is induced by it, is unclear.

Our pathway analysis also showed enrichment for other innate and adaptive immune related pathways in association with *Campylobacter* resistance. Of increased interest is the pathway related with IL-17 signalling since several previous studies suggested that IL-17 signalling and T_H_-17 responses play a role in resistance to *Campylobacter* colonisation in chickens following experimental inoculation [13, 16, 19, 56, 63, 64]. Future studies could seek to characterise the timing and magnitude of such responses in birds of predicted resistant or susceptible genotypes upon exposure. An important factor to be considered in future studies of this type is to characterise any concurrent infections and the subsequent relationship with *Campylobacter* and the host immune response. Pathway analysis also detected enrichment for the renin-angiotensin system, components of which have been reported to be activated by *Campylobacter* infection in chickens [16] and in the gastric mucosa of *Helicobacter pylori* infected humans [65].

## Conclusion

Our comprehensive genomic analyses estimated significant heritability of *Campylobacter* resistance in a commercial broiler population and identified QTLs, transcripts and networks in common with previous studies. A clear association with the MHC locus on chromosome 16 was identified, including detection of differentially transcribed MHC-related genes in the QTL interval in birds at the extreme of colonisation phenotype. The low frequency of susceptibility-associated alleles in the broiler population studied precluded the selection of predicted resistant or susceptible birds for experimental challenge. The advent of rapid genome-editing technology, for example reliant on modification of primordial germ cells implanted into sterile recipients during embryo development [66], provides a potential means to validate the role of genetic variation in *Campylobacter* resistance. The QTLs identified accounted for a c. 2 log_10_ CFU/g difference in caecal C. *jejuni* colonisation, sufficient to provide a significant reduction on the risk of contamination to human health [10]. However, resistance-associated variation was already highly prevalent in the population studied and environmental factors, which played a far greater role in the phenotype, may be more amenable to rapid and effective intervention. A multifactorial approach addressing both genetic and non-genetic factors is therefore needed to reduce *Campylobacter* levels in poultry and the incidence of the human disease attributed to this source.

## Materials and Methods

### Animals and sampling

A total of 3,000 birds of an outbred pure-bred commercial broiler line from the Aviagen breeding programme were housed within a non-bio-secure environment referred to as sib-test environment aimed to resemble broader commercial conditions and where full sibs and half sibs of selection candidates are placed [67]. Birds were fed standard maize-based starter, grower and finisher diets in line with industry practice. All birds throughout the study received the same vaccinations as per commercial regime and were reared under the same management practices and environmental parameters [26]. Birds were naturally exposed to *Campylobacter* spp. under these conditions, as confirmed by routine sampling of the environment using the ‘boot sock’ method described previously [25]. Birds were culled and phenotyped when they reached the age of five weeks. This was performed in batches of 100 birds (50 males and 50 females) giving a total of 3,000 birds phenotyped over a period of 16 months. After culling of birds by cervical dislocation by trained personnel, cardiac blood was collected for DNA extraction, the two caeca were collected for enumeration of viable *Campylobacter*, and the two caecal tonsils were stored in RNAlater (Thermo Fisher Scientific, Waltham, USA) for subsequent RNA extraction.

### Ethics statement

As these were commercial birds from an industry breeding programme and were not experimentally inoculated, the study was conducted outside the auspices of the Animals (Scientific Procedures) Act 1986, but was nevertheless subject to scrutiny and approval by the Animal Welfare & Ethical Review Body of The Roslin Institute, University of Edinburgh (under PPL 60/4420).

### Phenotyping and genotyping

To enumerate *Campylobacter*, serial ten-fold dilution series of weighed contents of the two caeca were separately prepared to 10^−7^ in phosphate-buffered saline and 100 μl of each dilution plated to modified charcoal deoxycholate (mCCDA) agar supplemented with cefoperazone (32 mg/L) and amphotericin B (10 mg/L; Oxoid), followed by incubation for 48 h under microaerophilic conditions (5% O_2_, 5% CO_2_, and 90% N_2_) at 41°C. Dilutions were plated in duplicate and colonies with morphology typical of *Campylobacter* were counted and expressed as CFU/g. The theoretical limit of detection by the method used was 100 CFU/g. In instances where no colonies were observed after direct plating, a *Campylobacter* load equal to the theoretical limit of detection was assumed, as enrichment to confirm the absence of *Campylobacter* in caecal contents was not performed.

All the birds were genotyped with a proprietary 50K high-density genome-wide SNP array and then imputed using AlphaImpute [68, 69] to the 600K SNP Affymetrix® Axiom® HD array [70] based on parent, grand-parent and great-grand-parent 600K SNP Affymetrix data. Of 3,000 birds sampled, genotypes for 2,718 birds were successfully imputed. Imputation failures likely reflect a lack of compatibility between the pedigree information and the genotypic data.

### Heritability analysis

As the distributions of *Campylobacter* levels were not normally distributed, all counts were log-transformed and expressed as log_10_ CFU/g. Genetic parameters were estimated for caecal *Campylobacter* colonisation resistance using a mixed linear univariate model that included the date of sampling and the sex as fixed effects, and the random effect of the individual birds linked to each other with the pedigree genetic relationship matrix. Body weight and maternal effects were also tested but their effects on the *Campylobacter* levels were not significant and therefore were not included in the final model. Genetic relationships between birds were calculated using a three generations pedigree and included in the analyses. The heritability of the trait was calculated as the ratio of the additive genetic variance to the total phenotypic variance. The analysis was performed using ASReml v4.0 [71].

### Genome-wide association study

The 50K and 600K SNP genotype data were subjected to quality control measures using PLINK v1.09 [72] with parameters of minor allele frequency >0.05, call rate >95 % and Hardy-Weinberg equilibrium (*P* > 10^−6^). After quality control, 37,498 and 288,381 SNP markers remained for further analysis (from the 50K and 600K datasets, respectively). Positions of SNP markers were obtained using the GalGal5 annotation, available via the Ensembl Genome Browser (www.ensembl.org). Population stratification was investigated using a genomic relatedness matrix generated from all individuals. This was converted to a distance matrix that was used to carry out classical multi-dimensional scaling analysis (MSA) using the R package GenABEL [73] to obtain its principal components. The GEMMA v0.98.1 algorithm [74] was used to perform GWAS analyses using a standard univariate linear mixed model in which date of sampling and sex were fitted as fixed effects and the genomic relatedness matrix among individuals was fitted as a polygenic random effect. After Bonferroni correction for multiple testing, significance thresholds for analysis with the 50K array were *P* ≤ 1.33 × 10^−6^ and *P* ≤ 2.66 × 10^−5^ for genome-wide significant levels (i.e., *P* ≤ 0.05) and suggestive significant levels (namely one false positive per genome scan), respectively, corresponding to -log_10_(*P*) of 5.87 and 4.47. The significance thresholds for the 600K array after Bonferroni correction were *P* ≤ 1.73 × 10^−7^ and *P* ≤ 3.46 × 10^−6^ corresponding to -log_10_(*P*) of 6.76 and 5.45. The extent of linkage disequilibrium (LD) between significant SNPs located on the same chromosome regions was calculated using the r-square statistic of PLINK v1.09 [72].

Individual markers found to be significant in the previous step were further examined with a mixed model that included the same fixed effects as used in the GWAS, the fixed effect of the corresponding SNP locus genotype and the random effect of the animal. Additive (a) and dominance (d) effects, and the proportion of total phenotypic variance (PV_P_) due to each SNP locus were calculated as follows:

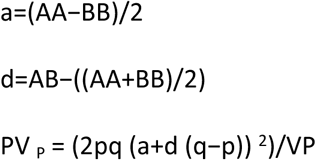

where AA, BB and AB were the predicted trait values of the respective genotypic classes, p and q were the allelic frequencies of A and B at the SNP locus, and VA and VP were the additive genetic and total phenotypic variance of the trait. The latter were estimated with the same model used for the heritability estimate. All analyses were run with ASReml v4.0 [71].

### Regional heritability mapping

RHM analyses were performed using DISSECT [75] fitting genomic regions of 20 SNPs in sliding windows along each chromosome with the same fixed effects as the ones used in the single SNP GWAS described above. The significance of genomic regions was assessed with the likelihood ratio test statistic, which was used to compare the RHM model where both the whole genome and a genomic region were fitted as random effects against a base model that excluded the latter effect. Only the 50K data was analysed with this method. A total of 1,915 regions were tested across the genome. After adjustment for multiple testing, using the Bonferroni correction, significance thresholds were *P* ≤ 2.63×10^−5^ and *P* ≤ 5.26×10^−5^ for genome-wide levels (*P* ≤ 0.05) and suggestive levels (namely one false positive per genome scan), respectively, corresponding to -log_10_(*P*) of 4.57 and 3.27.

### SNP and candidate region annotation

All significant SNPs identified in the GWAS and RHM analyses were mapped to the GalGal5 reference genome and annotated using the Ensembl Variant Effect Predictor (http://www.ensembl.org/Tools/VEP). Moreover, all the genes that were located within the 20 SNP windows found to be significant by RHM; and the 250 kb 5’ and 3’ regions of the significant SNP markers identified by the GWAS were annotated using GalGal5 data obtained by the BioMart data mining tool (http://www.ensembl.org/biomart/martview/). This allowed us to catalogue all the genes that were located in the vicinity of the identified significant SNP markers for *Campylobacter* colonisation resistance.

### Transcriptomic analyses

Total RNA was prepared from the caecal tonsils of 23 broilers, selected on their genotype (allele combination in the significant identified markers) and caecal *Campylobacter* load, after correction for other sources of systematic variation (sex and date of sampling). Details of the birds selected are shown in S3 Table. RNA was extracted using the RNeasy Mini Kit (Qiagen Hilden, Germany) according to manufacturer’s instructions. The resultant RNA was checked for quality using the Agilent Tapestation 2200, and all samples were of high quality with RNA Integrity Numbers (RIN) greater than 9. Library preparation was performed by Edinburgh Genomics (http://genomics.ed.ac.uk/) using the Illumina TruSeq mRNA (poly-A selected) library preparation protocol (Ilumina; Part: 15031047, Revision E). The mRNA was sequenced by Edinburgh Genomics at a depth of > 40 million strand-specific 75 bp paired-end reads per sample, using an Illumina HiSeq 4000. Expression levels for the 23 samples were estimated using Kallisto v0.43.0 [76]. Rather than aligning RNA-seq reads to a reference genome, reconstructing transcripts from these alignments and then quantifying expression as a function of the reads aligned, Kallisto employs a ‘lightweight’ algorithm, which first builds an index of k-mers from a known transcriptome. As a reference transcriptome, we obtained from Ensembl v89 the set of GalGal5 cDNAs and ncRNA transcripts (ftp://ftp.ensembl.org/pub/release-89/fasta/gallus_gallus/cds/Gallus_gallus.Gallus_gallus-5.0.cds.all.fa.gz, and ftp://ftp.ensembl.org/pub/release-87/fasta/gallus_gallus/ncrna/Gallus_gallus.Gallus_gallus-5.0.ncrna.fa.gz; n=38,118 total transcripts, representing 10,846 protein-coding genes and 937 non-coding genes). Expression levels were then estimated directly (i.e., in an alignment-free manner) by quantifying exact matches between reads and k-mers. Expression is reported per transcript as the number of transcripts per million, and is summarised to the gene level as described previously [77].

#### Differential expression

Differential expression analysis was run on caecal tonsils from birds with high, intermediate and low *Campylobacter* load, after adjusting for sex and seasonality, using the Kallisto output with the R/Bioconductor package ‘Sleuth’ v0.29.0 [78].Differential expression was considered significant for FDR corrected *P* values ≤ 0.05 and suggestive significant for uncorrected *P* values ≤ 0.001. Additional differential expression analyses were performed using the qRT-PCR output for a subset of the genes found to be significantly differentially expressed in the initial RNA-Seq analysis. Least square mean pairwise comparisons between different *Campylobacter* levels were conducted. Tukey’s HSD post-hoc test adjustment was applied at a significance level of 0.05.

#### Expression QTL analysis

eQTL analyses were performed using the R package Matrix eQTL v2.1.0 [79]. The number of transcripts per million, derived from Kallisto analysis as described above, were used as a measure of gene expression. Several covariates (log_10_ transformed *Campylobacter* counts, sex, date of sampling) were included in the association analysis. *Cis-* and *trans*-eQTLs were obtained, considering *cis-*acting SNPs to be within 100 kb from the 5’ start or 3’ end of a known gene. *P* values were corrected using false discovery rate (FDR) estimated with Benjamini-Hochberg procedure. SNP - gene expression association was considered significant for FDR corrected *P* values ≤ 0.05. For the significant *cis-*eQTL we estimated effect size using the log allelic fold-change (aFC) measurement. aFC is defined as the log-ratio between the expression of the haplotype carrying the alternative variant allele to the one carrying the reference allele and was calculated as described [80]. Briefly, the model assumes an additive model of expression in which the total expression of a gene in a given genotype group is the sum of the expression of the two haplotypes: e(genotype) = 2e_r, e_r + e_a, and 2e_a for reference homozygotes, heterozygotes, and alternate homozygotes, respectively, where e_r is the expression of the haplotype carrying the reference allele, and e_a the expression of the haplotype carrying the alternative allele. The allelic fold change k is defined as e_a = k e_r where 0 < k < ∞. aFC is represented on a log_2_ scale as s = log_2_ k.

#### SNP calling and allele-specific expression analysis

In order to perform allele-specific expression analysis we aligned RNA-Seq reads to the reference genome and called the genomic variance in the previously identified QTL regions. Quality filtering and removal of residual adaptor sequences from the raw reads was first performed using Trimmomatic v0.38 [81]. Leading and trailing bases with a Phred score less than 20 were removed, and the read trimmed if the average Phred score over a sliding window of four bases was less than 20. Only reads where both forward and reverse pairs were longer than 36 bp post-filtering were retained. Filtered reads were mapped to the chicken genome (Gallus_gallus-5.0; Genbank assembly GCA_000002315.3) [82] using STAR v2.6.1a [83], with the maximum number of mismatches allowed for each read pair set to 10% of the trimmed read length, and minimum and maximum intron lengths set to 20 bases and 1 Mb, respectively. PCR duplicates were marked and SNPs were identified and genotyped called for individual samples using samtools v1.6 [84], ignoring reads with mapping quality < 20 and bases with Phred quality scores < 20. SNPs within 5 bp of an indel, with mapping quality < 20, minor allele frequency (MAF) < 0.05 or where < 4 reads supported the alternative allele were also discarded. The SNPs located within the QTL regions identified by the GWAS were annotated using Variant Effect Predictor, as described above.

Allelic-specific expression was assessed using the R package AllelicImbalance v1.24.0 [85]. For every SNP in a region of interest, read counts were obtained for each allele present in a heterozygous animal, provided it was present in > 4 and < 17 heterozygous animals (i.e. 75% of the total animals). SNPs with less than 10 reads were excluded. A binomial test was performed to assess the significance of the difference in allelic count. Allele-specific expression was considered significant if the mean *P* value across all heterozygotes was ≤ 0.05.

### Quantitative RT-PCR validation of differentially expressed genes

First strand synthesis was performed using 1 μg of total RNA and the Verso cDNA Synthesis Kit (Thermo Scientific) according to the manufacturer’s instructions. qPCR reactions were performed using the Forget-Me-Not™ qPCR Master Mix (Biotium) in 20 μL volumes containing 1 X Forget-Me-Not™ qPCR Master Mix, 0.5 μM of each forward and reverse primer, 50 nM of ROX reference dye and 2 μL of cDNA at a 1:10 dilution in template buffer. Gene-specific primers were designed and purchased from Sigma. Primer sequences are detailed in S10 Table. The amplification and detection of specific DNA was achieved using the AB 7500 FAST Real-Time PCR System (Applied Biosystems) and the following programme: 95°C for 2 min followed by 40 cycles of 95°C for 5 s then 60°C for 30 s. To confirm the presence of a single PCR product, melting curves were generated by one cycle of 60°C for 1 min, increasing to 95°C in 1% increments every 15 s. Samples were run in triplicate and each qPCR experiment contained 3 no-template control wells and a 5-fold dilution series in duplicate of pooled caecal tonsil derived cDNA from several birds from which standard curves were generated. The expression of genes were normalised to the geometric mean of three reference genes found previously to be stably expressed in chicken lymphoid organs; *r28S, TBP* and *GAPDH* [86].

### Pathway, network and functional enrichment analyses

Identification of potential canonical pathways and networks underlying the candidate genomic regions associated with *Campylobacter* colonisation resistance was performed using the Ingenuity Pathway Analysis (IPA) programme (www.ingenuity.com). IPA constructs multiple possible upstream regulators, pathways and networks that serve as hypotheses for the biological mechanism underlying the phenotypes based on a large-scale causal network derived from the Ingenuity Knowledge Base. IPA then infers the most suitable pathways and networks based on their statistical significance, after correcting for a baseline threshold [87]. The IPA score in the constructed networks can be used to rank these networks based on the P values obtained using Fisher’s exact test (IPA score or *P* score = –log_10_(*P* value)).

The gene list for *Campylobacter* colonisation resistance was also analysed using the Database for Annotation, Visualization and Integrated Discovery (DAVID) v6.8 [88]. Gene ontology (GO) was determined and functional annotation clustering analysis was performed. The *Gallus gallus* background information is available in DAVID and was used for the analysis. The enrichment score (ES) of the DAVID package is a modified Fisher exact *P* value calculated by the software, with higher ES reflecting more enriched clusters. An ES greater than 1 means that the functional category is overrepresented.

## Acknowledgements

The authors gratefully acknowledge the support of the Biotechnology & Biological Sciences Research Council via the LINK scheme (grant reference BB/J006815/1) and Institute Strategic Programme funding at The Roslin Institute (BBS/E/D/20231760 and BBS/E/D/20002172). We also acknowledge funding from the Scottish Government via the Rural & Environmental Science and Analytical Services programme of research for 2016-2021. These funders had no role in study design, data collection and analysis, decision to publish, or preparation of the manuscript. We dedicate this manuscript to our late colleagues Dr Paul Hocking and Professor Pete Kaiser, who played key roles in conception of the study and supervision of the research.

## Author contributions

PK, KW, MS, PH, MF, AK, SA, RB and AP conceived and designed the genetic study of *Campylobacter* resistance and secured substantial funding; MS, AP and DH conceived and designed the transcriptomic study of *Campylobacter* resistance and MS and AP secured substantial funding. AP, LR, RB and KW performed data collection. AP and LR performed the phenotyping, with input from MS. AK performed the imputation of the genomic data. AP and GB, with input from AK, PK and KW, collated and edited the genotyping data and performed the genetic and genomic analysis. AP performed the transcriptomic analyses with input from KR, SB, DH and MS. DR performed the ALE analysis. AP performed the pathway analyses. KW reconstructed the MHC haplotypes. AP performed the haplotype analyses and interpreted the results with input from JK and KW. KR performed the RNA extractions, KR and AB performed the qRT-PCR to validate the RNA-Seq data. AP, MS, KW, AK, GB, DH, JK, MF, RB and SA interpreted these results. AP and MS wrote the manuscript. All other co-authors provided manuscript editing and feedback. All authors read and approved the final manuscript.

## Availability of data

The sequencing and expression data from caecal tonsils of chickens with different levels of *Campylobacter* colonisation in their caeca are deposited in the European Nucleotide Archive under accession number PRJEB22580.

---

**Fig. 6. Pathway analysis using the IPA software.** The most highly represented canonical pathways derived from genes located within the candidate regions for *Campylobacter* colonisation resistance in commercial chickens. The solid yellow line represents the significance threshold. The line joining squares represents the ratio of the genes represented within each pathway to the total number of genes in the pathway.

## Supporting Information

**S1 Fig. Patterns of linkage disequilibrium (LD) for SNP markers on chromosome 16 associated significantly with *Campylobacter* colonisation resistance in commercial chickens**. All the significant SNP markers were in high LD, illustrated with red colour, and were located within the same LD block (230kb) marked with triangle.

**S2 Fig. Manhattan plot and Q-Q plot displaying the RHM results for chicken *Campylobacter* colonisation resistance**. (A) Genomic location is plotted against –log_10_(P) in the Manhattan plot. Genome-wide (*P* < 0.05) and suggestive genome-wide thresholds are shown as red and blue lines, respectively. (B) Q–Q plot of observed *P* values against the expected *P* values for *Campylobacter* caecal load (log-transformed CFU of *Campylobacter* per gram of caecal content).

**S3 Fig. Pathway analysis using the IPA software**. The most highly represented canonical pathways derived from differentially expressed genes in the caecal tonsils of commercial chickens with divergent caecal *Campylobacter* load. The solid yellow line represents the significance threshold.

**S4 Fig. Network analysis results using the IPA software**. A gene network related with cell death and survival and organismal injuries and abnormalities were constructed from differentially expressed genes in the caecal tonsils of commercial chickens with divergent caecal *Campylobacter* load. Arrows with solid lines represent direct interactions and arrows with broken lines represent indirect interactions. Genes with red labels are upregulated, with green labels downregulated and with white labels are those added to the IPA analysis because of their interaction with the target gene products.

**S1 Table. List of SNP windows associated with *Campylobacter* colonisation resistance in Regional Heritability Mapping (RHM) analysis of the commercial chickens**.

**S2 Table. List of genes and non-coding RNAs located in the candidate regions for *Campylobacter* resistance identified by GWAS and RHM analysis in commercial chickens**.

**S3 Table. Details of the commercial chickens with low, average and high *Campylobacter* colonisation levels selected for RNA-Seq**.

**S4 Table. Expression (cis-) QTL analysis results**. List of SNPs identified by GWAS which were acting as *cis-*elements for genes located in the candidate regions for *Campylobacter* colonisation resistance.

**S5 Table. Expression (trans-) QTL analysis results**. List of SNPs identified by GWAS which were acting as *trans-*elements across the chicken genome.

**S6 Table. Allele specific expression (ASE) analysis results**. List of SNPs in the candidate regions for *Campylobacter* colonisation resistance which had a significant ASE.

**S7 Table. Functional annotation clustering analysis of the genes located in the candidate regions for *Campylobacter* colonisation resistance in chickens**.

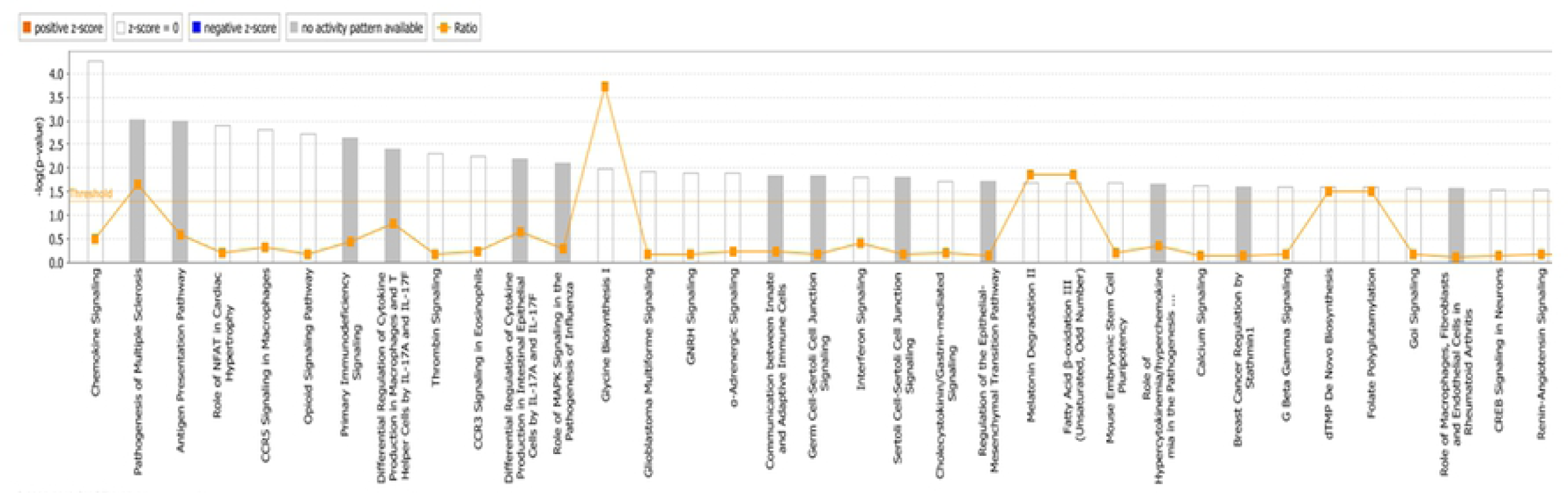

